# SARS-CoV-2 Nsp14 activates NF-κB signaling and induces IL-8 upregulation

**DOI:** 10.1101/2021.05.26.445787

**Authors:** Taiwei Li, Adam D. Kenney, Helu Liu, Guillaume N. Fiches, Dawei Zhou, Ayan Biswas, Jianwen Que, Netty Santoso, Jacob S. Yount, Jian Zhu

**Affiliations:** Department of Pathology, The Ohio State University Wexner Medical Center, Columbus, OH 43210, USA; Department of Microbial Infection and Immunity, The Ohio State University Wexner Medical Center, Columbus, OH 43210, USA; Department of Medicine, Columbia University Medical Center, New York, NY 10032, USA; Department of Genetics, The University of Alabama at Birmingham, Birmingham, AL 35233, USA

**Keywords:** SARS-CoV-2, NF-κB, IL-8, IMPDH2, ribavirin, mycophenolic acid

## Abstract

Severe acute respiratory syndrome coronavirus 2 (SARS-CoV-2) infection leads to NF-κB activation and induction of pro-inflammatory cytokines, though the underlying mechanism for this activation is not fully understood. Our results reveal that the SARS-CoV-2 Nsp14 protein contributes to the viral activation of NF-κB signaling. Nsp14 caused the nuclear translocation of NF-κB p65. Nsp14 induced the upregulation of IL-6 and IL-8, which also occurred in SARS-CoV-2 infected cells. IL-8 upregulation was further confirmed in lung tissue samples from COVID-19 patients. A previous proteomic screen identified the putative interaction of Nsp14 with host Inosine-5’-monophosphate dehydrogenase 2 (IMPDH2) protein, which is known to regulate NF-κB signaling. We confirmed the Nsp14-IMPDH2 protein interaction and found that IMPDH2 knockdown or chemical inhibition using ribavirin (RIB) and mycophenolic acid (MPA) abolishes Nsp14-mediated NF-κB activation and cytokine induction. Furthermore, IMDPH2 inhibitors (RIB, MPA) efficiently blocked SARS-CoV-2 infection, indicating that IMDPH2, and possibly NF-κB signaling, is beneficial to viral replication. Overall, our results identify a novel role of SARS-CoV-2 Nsp14 in causing the activation of NF-κB.

## Introduction

SARS-CoV-2 is a beta-coronavirus that causes the current, severe COVID-19 pandemic globally. The viral genome of SARS-CoV-2 is a ~30 kb polycistronic, positive-strand RNA that encodes multiple structural and nonstructural proteins *(1, 2)*. SARS-CoV-2 nonstructural proteins (Nsp1-16) play diversified roles in supporting viral RNA/protein synthesis and virion assembly, including manipulating host gene expression and host antiviral responses *(3, 4)*. It has been recently reported that SARS-CoV-2 infection suppresses type I interferon (IFN) signaling *(5, 6)*, while it induces the activation of NF-κB signaling that plays a central role in the production of pro-inflammatory cytokines, including interleukin (IL)-6 and IL-8 *(5, 7, 8)*. In certain cases, massive inflammatory responses occur due to hyper-activation of the immune system, resulting in a widespread and uncontrolled cytokine storm, leading to acute respiratory distress syndrome (ARDS), life-threatening lung damage, and increased mortality of COVID-19 patients. However, the underlying mechanism of how SARS-CoV-2 infection contributes to NF-κB-mediated inflammatory responses that are expected to determine the outcome of SARS-CoV-2 viral replication and pathogenesis is still largely uncharacterized.

Here we focused on characterizing the regulatory functions of SARS-CoV-2 Nsp14 that are required for efficient viral replication. Nsp14 is a conserved, multifunctional viral factor participating in synthesizing and modifying coronaviral sub-genomic (sg) RNAs *(9)*. Nsp14 possesses a 3’ to 5’ exonuclease activity that excises mismatched base pairs during viral RNA replication *(10-12)*, providing a proofreading function that increases the fidelity of viral RNA synthesis *(13, 14)*. Nsp14 also possesses RNA methyltransferase activity required for guanine-N7 methylation *(15)*. Nsp14-mediated guanine-N7 methylation cooperates with 2’-O RNA methylation mainly catalyzed by Nsp10/16, leading to 5’-capping of newly synthesized sgRNAs *(16, 17)*, which not only prevents degradation by host RNA 5’ exonucleases and recognition by host foreign RNA sensors, such as RIG-I *(18)*, but also increases translation efficiently of host ribosomes to synthesize viral proteins *(19, 20)*. Nsp14 has also been reported to reduce the accumulation of viral double-stranded (ds) RNAs and thus dampen the pathogen-associated molecular pattern (PAMP) mediated antiviral response *(21)*. In addition, Nsp14 is known to facilitate recombination between different viral RNAs to generate new strains *(22)*. Compared to these well-studied viral functions of Nsp14, its regulation of host cellular events is much less investigated. An earlier large-scale proteomic analysis reporting candidate interacting partners for all of the SARS-CoV-2 open reading frames (ORFs) indicated that the host inosine-5’-monophosphate dehydrogenase 2 (IMPDH2) protein is one binding partner of SARS-CoV-2 Nsp14 protein *(23)*. Interestingly, IMPDH2 has been identified to play a role in regulating NF-κB signaling *(24)*. Our new results showed that SARS-CoV-2 Nsp14 activates NF-κB signaling and induces IL-8 upregulation, which indeed requires the interaction of Nsp14 with IMPDH2.

## Results

### SARS-CoV-2 Nsp14 causes activation of NF-κB

We initially investigated the effect of SARS-CoV-2 Nsp14 along with Nsp10 and Nsp16 on certain immune signaling pathways. The pcDNA-V5-FLAG-Nsp14/10/16 vectors were individually transfected in HEK293T, and the expression of the individual proteins was confirmed (**Fig S1A**). We then utilized these expression vectors for interferon-sensitive response element (ISRE) and NF-κB luciferase reporter assays (**Fig S1B** and **C**). Nsp14 mildly increased ISRE activity at the basal level but caused its decrease in IFN-α-treated HEK293T cells, while Nsp10 and Nsp16 mildly decreased ISRE activity at both conditions, which is consistent with earlier findings *(3, 4)*. On the contrary, only Nsp14 significantly increased NF-κB activity in both untreated and TNF-α-treated HEK293T cells. TNF-α did not affect the expression of transfected Nsp14 in HEK293T cells (**Fig 1A**) but induced a drastic increase of NF-κB activity that was further enhanced by Nsp14 (**Fig 1B**). Thus, we further investigated Nsp14-induced activation of NF-κB signaling. The impact of Nsp14 on nuclear localization of NF-κB p65 was determined in HEK293T cells transfected with Nsp14. Indeed, Nsp14 expression led to the significant increase of nuclear but not total p65 protein (**Fig 1C, D** and **Fig S2**). These results confirmed that SARS-CoV-2 Nsp14 activates NF-κB signaling.

**Fig 1.**
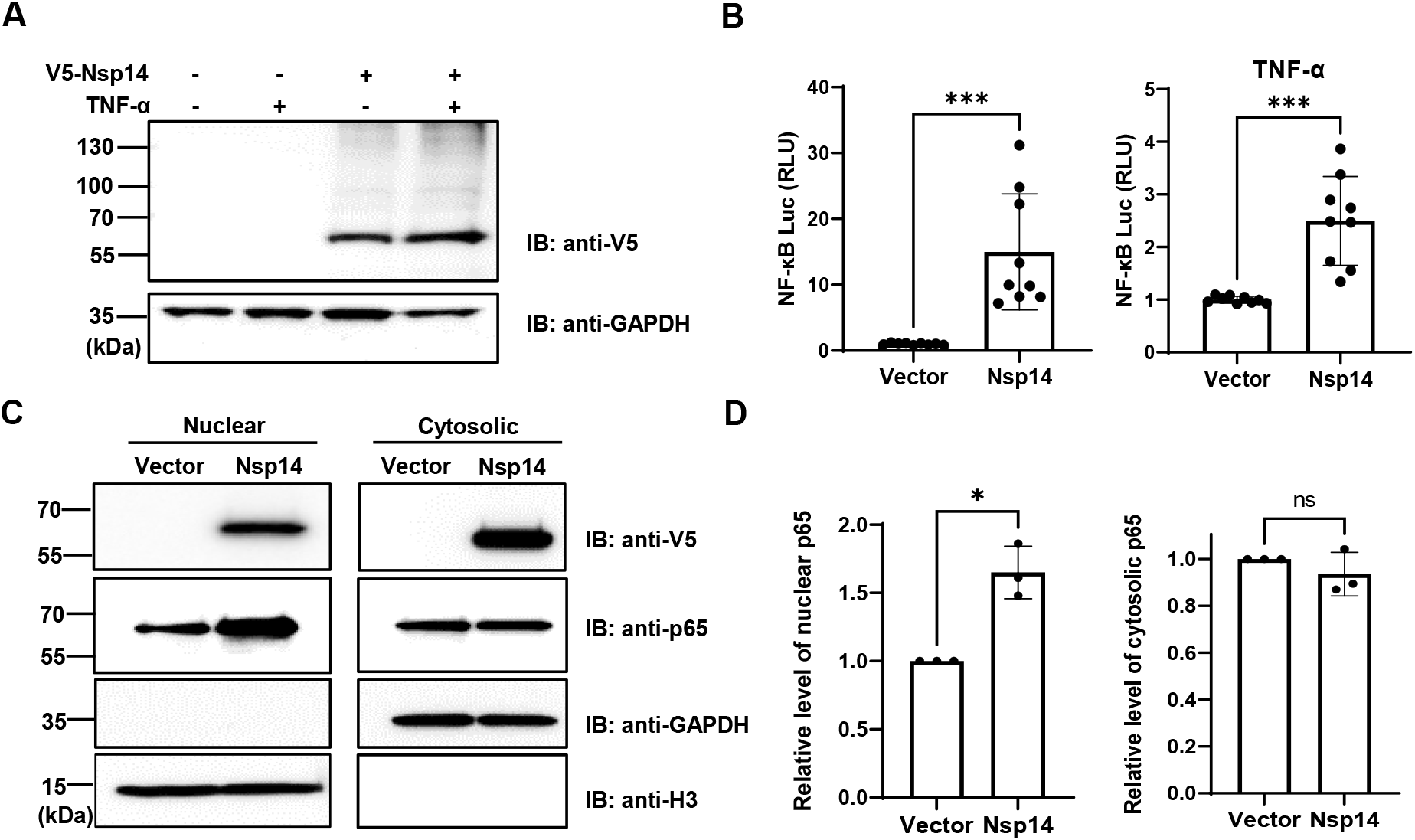
SARS-CoV-2 Nsp14 increases NF-κB activity. **(A-C)** HEK293T cells were transiently transfected with V5-FLAG-Nsp14 or empty vector, and treated with or without TNF-α. V5-FLAG-Nsp14 was analyzed by protein immunoblotting (**A**). HEK293T cells transfected with V5-FLAG-Nsp14 or empty vector along with NF-κB-driven firefly luciferase and TK-driven renilla luciferase reporter vectors were un-treated or treated with TNF-α (**B)**. Luciferase activity (firefly/renilla) in these cells was measured and normalized to the empty vector. HEK293T cells transfected with V5-FLAG-Nsp14 or empty vector were subjected to the nuclear/cytosolic fractionation. V5-FLAG-Nsp14 and NF-κB p65 in the nucleus or cytosol were analyzed by protein immunoblotting (**C**). Histone H3 was used as the nuclear marker. The intensity of the p65 protein band was quantified and normalized to the empty vector (**D**). Results were calculated from 3 independent experiments and presented as mean +/- standard deviation (SD). (* p <0.05; *** p <0.001 by unpaired Student’s t-test).

### SARS-CoV-2 Nsp14 induces upregulation of IL-8

NF-κB plays a critical role in regulating pro-inflammatory gene expression. Since we showed that Nsp14 causes NF-κB activation, we further determined whether Nsp14 induces the expression of several interleukins (IL-4, 6, 8). IL-6 and IL-8 are defined gene targets of NF-κB *(25-27)*. In HEK293T cells transfected with pcDNA-V5-FLAG-Nsp14, IL-6 and IL-8 were consistently and significantly upregulated with or without TNF-α (**Fig 2A**). Results were similar in Nsp14-transfected A549 cells, although it was significant only in experiments without TNF-α (**Fig 2B**). As a control, we confirmed that TNF-α does not affect the expression of transfected Nsp14 in A549 cells (**Fig S3**). In contrast to IL-6 and Il-8, IL-4 was not induced by Nsp14 in HEK293T nor A549 cells.

**Fig 2.**
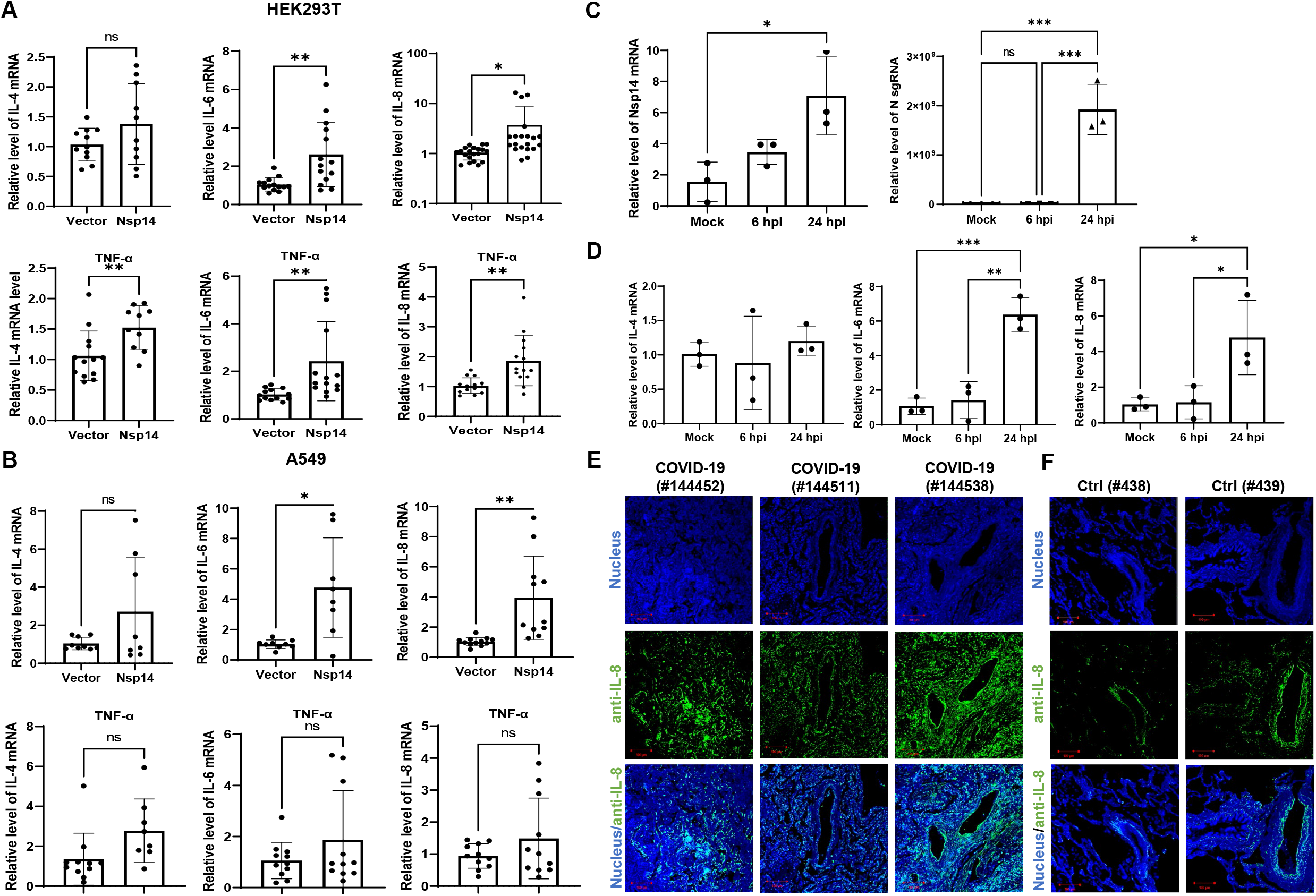
SARS-CoV-2 Nsp14 increases IL-6/8 expression. **(A)** HEK293T cells were transfected with V5-FLAG-Nsp14 or empty vector were un-treated or treated with TNF-α. The mRNA level of IL-4/6/8 in these cells was measured and normalized to the empty vector. **(B)** A549 cells were treated similarly as in (**A**) and analyzed for IL-4/6/8 expression. Results were calculated from at least 3 independent experiments and presented as mean +/- standard deviation (SD). (* p <0.05; ** p <0.01; by unpaired Student’s t-test). **(C, D)** HEK293T-ACE2 cells were infected with wild-type SARS-Cov-2 viruses. Cells were harvested at the indicated time points. Total RNAs were extracted, and expression of viral genes (Nsp14, N-protein, **C**) or ILs (IL-4, 6, 8, **D**) was analyzed by RT-qPCR and normalized to mock infection. Results were calculated from 3 technical repeats and presented as mean +/- standard deviation (SD). (* p <0.05; ** p <0.01; *** p <0.001 by one-way ANOVA and Tukey’s multiple comparison test). **(E, F)** Dissected lung tissues from COVID19 patients (**E**, donors #144452, #144511, #144538) or non-infected donors (**F**, donors #438, #439) were analyzed for IL-8 expression by immunofluorescence (green). Nuclei were stained with Hoechst (blue). Scale bar: 100 µm.

We next confirmed whether infection of cells with SARS-CoV-2 also induces upregulation of IL-6 and IL-8. HEK293T-ACE2 cells were infected with the SARS-CoV-2 viral strain USA-WA1/2020 *(28)*. Expression of viral genes, Nsp14 and nucleocapsid [N], was readily detected (**Fig 2C**). The SARS-CoV-2 infection also led to the upregulation of IL-6 and IL-8, but not IL-4 (**Fig 2D**). We employed immunofluorescence staining assays to determine whether IL-8 upregulation occurres in lung tissue samples dissected from deceased COVID-19 patients. The results showed that IL-8 expression is consistently higher in COVID-19 patients (**Fig 2E**) compared to un-infected cases (**Fig 2F**). IL-6 induction in the lung of COVID-19 patients has already been reported elsewhere *(29, 30)*. We primarily focused on IL-8 as the representative target gene of NF-κB for further analysis since its induction by Nsp14 is overall more robust than IL-6.

### IMPDH2 binds to Nsp14 and contributes to Nsp14 induction of IL-8

We first confirmed the putative protein interaction of Nsp14 with IMPDH2 *(23)* by protein co-immunoprecipitation (co-IP) assays in HEK293T cells co-transfected with the pLEX-V5-IMPDH2 and pEZY-FLAG-Nsp14 vectors (**Fig 3A**). As the next step, we determined whether endogenous IMPDH2 is required for IL-8 induction by Nsp14. IMPDH2-targeting or non-targeting (NT) siRNAs were transfected in HEK293T cells, and efficient knockdown of endogenous IMPDH2 was confirmed (**Fig 3B**). Remarkably, IMPDH2 knockdown abolished the IL-8 induction by Nsp14 in HEK293T cells without or with TNF-α (**Fig 3C**). However, overexpression of IMPDH2 had no significant effect on NF-κB activation by Nsp14 in HEK293T cells with or without TNF-α (**Fig S4**).

**Fig 3.**
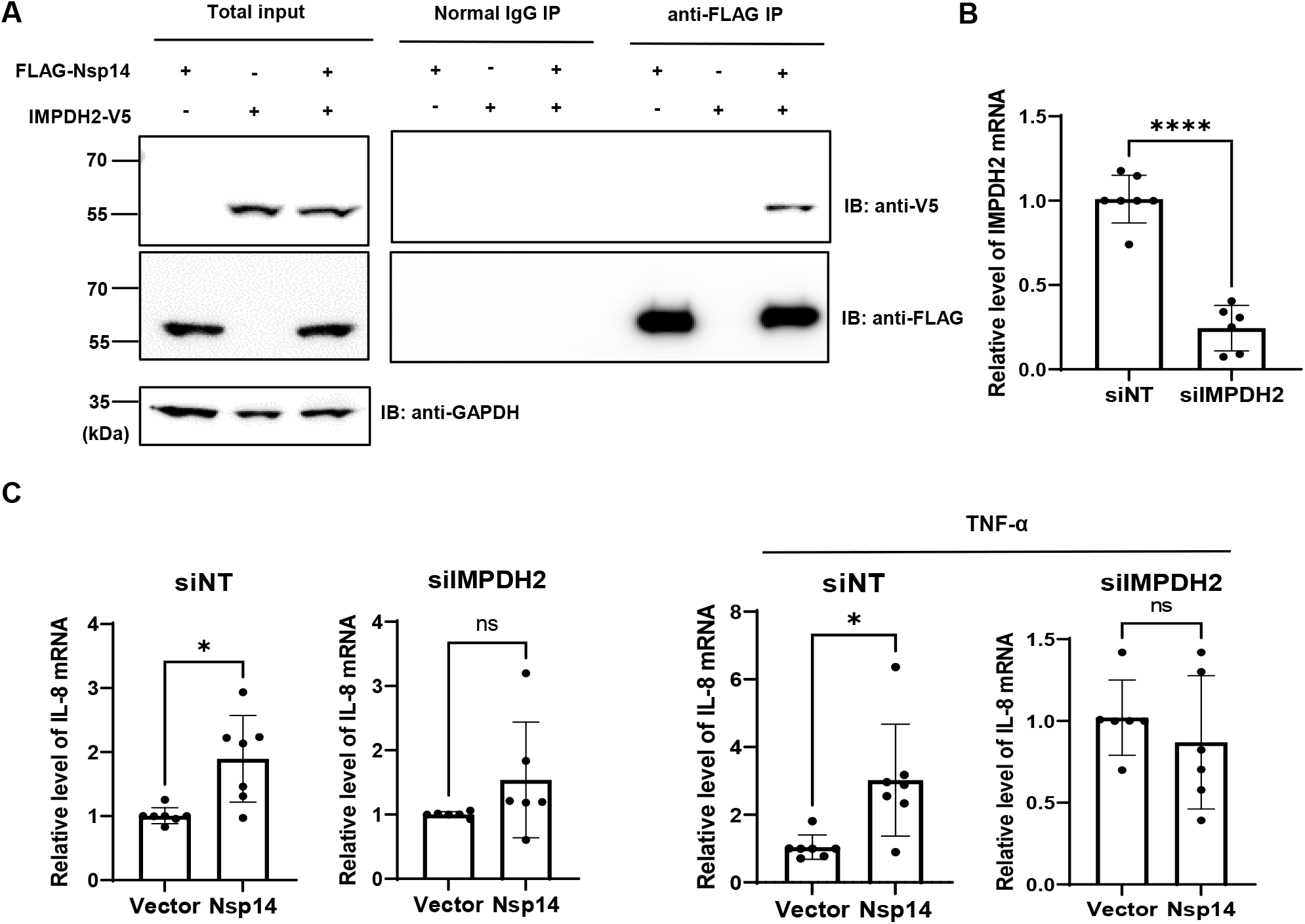
IMPDH2 associates with Nsp14 and is required for IL-8 upregulation by Nsp14. **(A)** HEK293T cells were transiently transfected with the vector expressing FLAG-Nsp14 or V5-IMPDH2, alone or together. Cell lysates were prepared and subjected to protein co-immunoprecipitation (co-IP) assays using anti-FLAG or control IgG antibody. Precipitated protein samples were analyzed by protein immunoblotting using anti-V5 and anti-FLAG antibodies. **(B)** HEK293T cells were transiently transfected with IMPDH2 or non-targeting (NT) siRNAs. mRNA level of IMPDH2 was measured and normalized to siNT. (**C**) HEK293T cells transfected with IMPDH2 or NT siRNAs were further transfected with V5-FLAG-Nsp14 or empty vector. These cells were untreated or treated with TNF-α. Total RNAs were extracted. IL-8 mRNA was analyzed and normalized to the empty vector. Results were calculated from 3 independent experiments and presented as mean +/- standard deviation (SD). (* p <0.05; **** p <0.0001 by unpaired Student’s t-test).

### IMPDH2 inhibition blocks Nsp14-mediated NF-κB activation and IL-8 induction

Since IMPDH2 is required for IL-8 induction by Nsp14, we expected that its inhibition would reduce Nsp14-mediated NF-κB activation and IL-8 induction. We tested two reported IMPDH2 inhibitors, ribavirin (RIB) and mycophenolic acid (MPA) *(23, 31)*. RIB is a synthetic nucleoside that occupies the IMPDH2 catalytic site to inhibit IMP conversion to xanthosine 5’-phosphate (XMP) during the guanine nucleotide (GTP) biosynthesis *(31-33)*. MPA shares similar features with the IMPDH2 cofactor, nicotinamide adenine dinucleotide (NAD^+^). MPA stacks and traps the XMP intermediate at the catalytic site to inhibit IMPDH2 enzyme activity *(31, 34)*. We confirmed that NF-κB activation by Nsp14 significantly decreases in HEK293T cells treated with RIB (**Fig 4A**) or MPA (**Fig 4B**) at multiple doses in the absence or presence of TNF-α using the NF-κB luciferase reporter assays. Likewise, treatment of HEK293T cells with RIB (**Fig 4C**) or MPA (**Fig 4D**) also caused the reduction of IL-8 induction by Nsp14. We next tested whether IMPDH2 inhibitors (RIB, MPA) also repress SARS-CoV-2 infection *in vitro*, considering that virus-mediated NF-κB activation would likely benefit its replication *(35-38)*. Indeed, we showed that the infection rate of SARS-CoV-2 decreases in both A549-ACE2 and HEK293T-ACE2 cells treated with RIB or MPA through quantification of cells expressing N protein by immunofluorescence staining assays (**Fig 4E-F, S5A-B**) or sgRNA level by RT-qPCR (**Fig 4G, S5C**). Consistently, we also identified that treatment of RIB or MPA leads to a significant reduction of IL-8 expression (**Fig 4H**).

**Fig 4.**
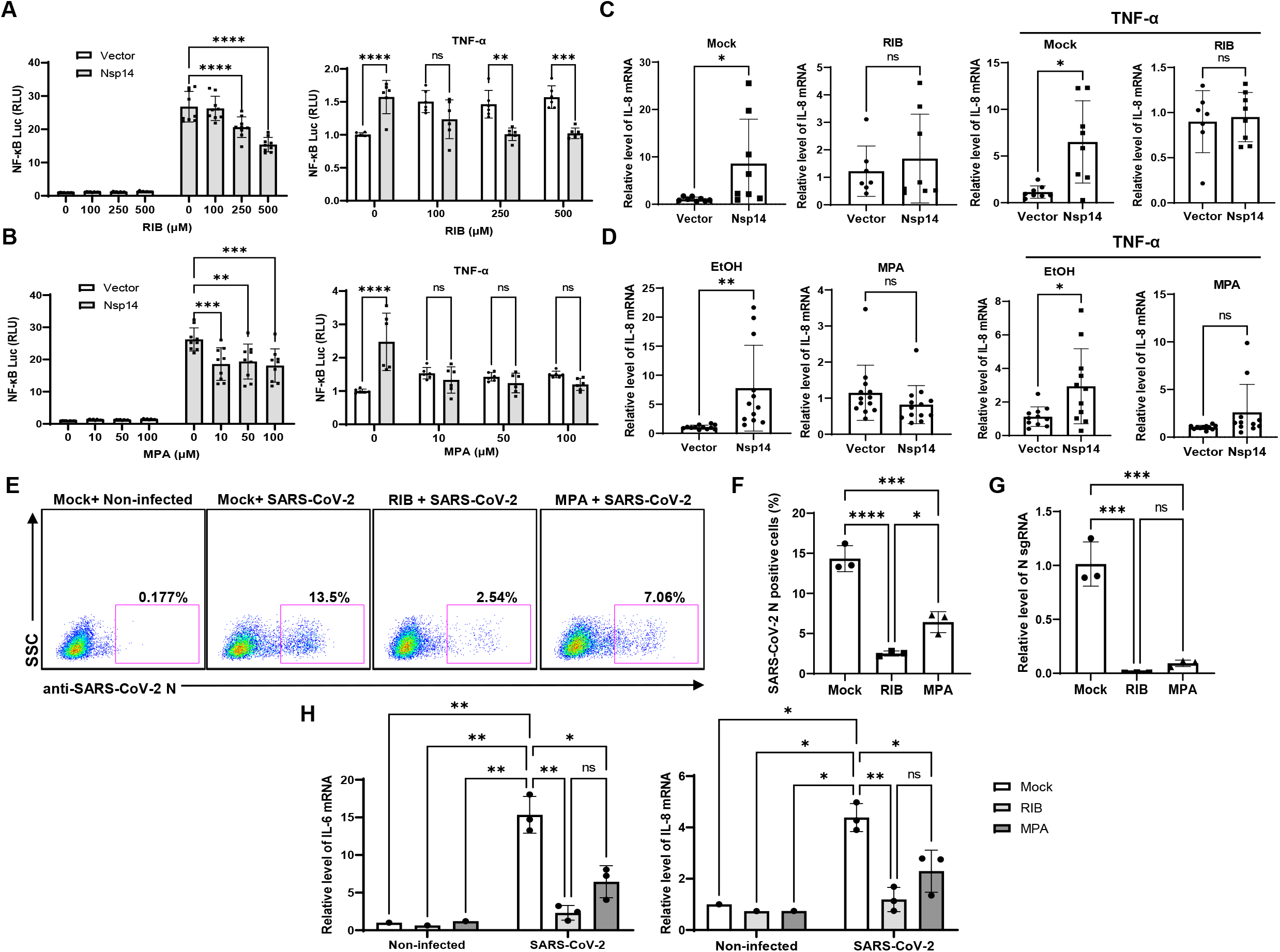
IMPDH2 inhibition reduces Nsp14-mediated NF-κB activation and IL8 induction. **(A)** HEK293T cells transfected with V5-FLAG-Nsp14 or empty vector along with NF-κB-driven firefly luciferase and TK-driven renilla luciferase reporter vectors were treated with ribavirin (RIB) at the basal or TNFα-stimulated condition. Luciferase activity (firefly/renilla) in these cells was measured and normalized to that of un-treated, empty vector-transfected cells. **(B)** Mycophenolic acid (MPA) was tested similarly as in (**A**). Results were calculated from at least 2 independent experiments and presented as mean +/- standard deviation (SD). (** p < 0.01; *** p < 0.001; **** p < 0.0001 by two-way ANOVA and Tukey’s multiple comparison test). **(C)** HEK293T cells transfected with V5-FLAG-Nsp14 or empty vector were treated with RIB at the basal or TNF-α-stimulated condition. Total RNAs were extracted. IL-8 mRNA was analyzed and normalized to the mock treatment. **(D)** MPA was tested similarly as in (**C**), and results were normalized to the solvent control (0.1% ethanol, EtOH). Results were calculated from 3 independent experiments and presented as mean +/- standard deviation (SD). (* p <0.05 by unpaired Student’s t-test). (**E-H**) A549-ACE2 cells were treated with RIB (500 µM), MPA (100 µM), or mock, and infected with SARS-Cov-2 viruses for 24 h. The SARS-CoV-2 infection was detected by intracellular staining of SARS-CoV-2 N protein (**E**). Percentage of SARS-CoV-2 N protein positive cells was calculated (**F**). Cells were harvested for RNA extraction, and N protein sgRNA was analyzed and normalized to the mock treatment (**G**). mRNA of IL-6 and IL-8 was analyzed and normalized to the non-infected cell with the mock treatment (**H**). Results were calculated from 3 technical repeats and presented as mean +/- standard deviation (SD). (* p <0.05; ** p <0.01; *** p <0.001; **** p <0.001 by two-way ANOVA and Tukey’s multiple comparison test).

## Discussion

Besides the well-known viral functions of SARS-CoV-2 Nsp14 to control modification and replication of viral RNA genomes, earlier studies illustrated that Nsp14 suppresses Type 1 IFN signaling and nuclear translocation of IRF3 to facilitate viral invasion of the host’s antiviral immune response *(3, 4)*. Our results showed that Nsp14, which is expressed at the early stage of primary infection *(7)*, also affects other cell signaling pathways, such as NF-κB signaling (**Fig 1**), likely to support viral replication. Activation of NF-κB may further trigger the production of downstream pro-inflammatory cytokines to initiate the cytokine storm and contribute to ARDS. In this study, we identified that Nsp14 increases nuclear translocation of p65 and induces expression of NF-κB’s downstream cytokines, such as IL-6 and IL-8, which have also been detected in lung tissues of COVID-19 patients *(5, 29)* and animal models of SARS-CoV-2 infection *(7)*. These cytokines are reported to play a critical role in regulating the recruitment and infiltration of immune cells (macrophages, neutrophils) during viral infection *(39, 40)*. Infiltrating immune cells may further escalate inflammatory responses leading to lung damage. Indeed, we showed that IL-8 expression is much higher in lung tissue samples of COVID-19 patients than in uninfected controls (**Fig 2E, F**).

Another key finding is that IMPDH2 is a host mediator of Nsp14 involved in NF-κB activation, verified by both genetic knockdown (**Fig 3**) and chemical inhibition (**Fig 4**). We confirmed the protein interaction of Nsp14 with IMPDH2, which was initially reported in earlier proteomic studies *(23, 29)*. Previous results also suggested that IMPDH2 benefits budding of Junín mammarenavirus (JUNV), propagation of lymphocytic choriomeningitis virus (LCMV) *(41)*, and replication of human norovirus (HuNV) *(42)*. IMPDH2 inhibitors have been used for treating hepatitis C virus (HCV) *(31, 43)*. Our results suggested that IMPDH2 likely supports the SARS-CoV-2 infection and Nsp14-mediated NF-κB activation as well. IMPDH2 is a protein target of certain immunosuppressive drugs used for organ transplantation and allograft rejection *(34, 44, 45)*, and it has been reported to regulate NF-κB signaling *(24, 46)*. Nsp14 may hijack IMPDH2 for NF-κB activation *(24)*, contributing to abnormal inflammatory responses. In terms of possible molecular mechanisms, since IMPDH2 participates in regulating the host nucleotide metabolism *(47, 48)*, it may further modulate cellular stress response and downstream NF-κB activation *(48-50)*. Nsp14 may manipulate IMPDH2 to increase the phosphorylation of IKKβ and IκBα to promote nuclear translocation and phosphorylation of p65 *(24)*. In addition, we also noticed that Nsp14 partially localizes in the nuclei of cells (**Fig 1C, D**), similar to findings from other groups *(51, 52)*. Thus, Nsp14 may associate with and modify the host cellular RNAs via its exonuclease and methyltransferase activities. Nsp14 may also affect the transcriptional activity of nuclear p65 and the expression of its gene targets. Future studies will be needed for further understanding how Nsp14 and IMPDH2 cooperate to activate NF-κB.

Our study has the translational significance since we showed that IMPDH2 inhibitors, RIB and MPA, effectively reduce viral replication of SARS-CoV-2 and expression of NF-κB’s downstream cytokines (IL-6 and IL-8) induced by SARS-CoV-2 (**Fig 4E-H**). It has been reported that IL-8 increases the replication of human immunodeficiency virus-1 (HIV-1), HCV, and cytomegalovirus (CMV) *(53-56)*. SARS-CoV-2 Nsp14 induces the NF-κB signaling and downstream cytokines, which may support the host cell proliferation and survival, or prevent cell apoptosis, thus benefiting viral replication *(38, 57)*. RIB and MPA are both FDA-approved drugs for treating HCV infection and transplant organ rejection, respectively. Our findings are supported by recent results showcasing the therapeutic potential of RIB and MPA for treating COVID-19 and SARS-CoV-2 infection. The combination of RIB with IFN β-1b and Lopinavir–Ritonavir therapy is currently in clinical trials for treating SARS-CoV-2 infection *(58)*, which has been shown to significantly alleviate the COVID-19 symptoms and suppress IL-6 levels in serum. In another preclinical study, MPA was reported to inhibit SARS-CoV-2 replication *(59)* and viral entry *(60)*. Our study delineated a potentially new mode of action (MOA) for these IMPDH2 inhibitors, which may disrupt the Nsp14-IMPDH2 axis that plays a crucial role in regulating activation of NF-κB signaling and induction of its downstream cytokines (**Fig 5**).

**Fig 5.**
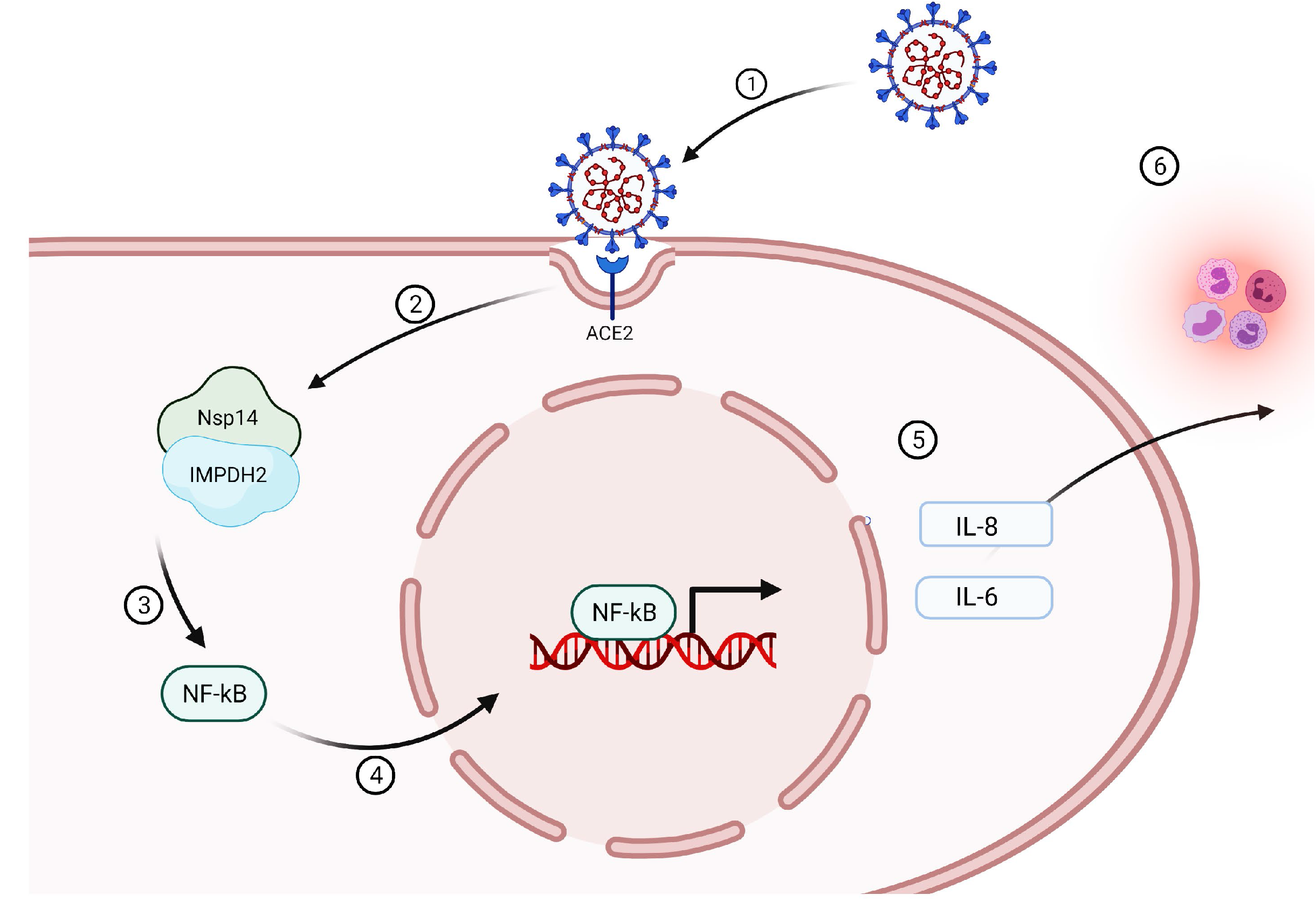
A working model of Nsp14-mediated NF-κB activation in SARS-CoV-2 infection. Infection of SARS-CoV-2 (**1**) leads to the expression of Nsp14 (**2**) that interacts with IMPDH2 (**3**). Such interaction promotes the nuclear translocation of NF-κB p65 (**4**) and its activation, which upregulates the expression of downstream cytokines, including IL-6 and IL-8 (**5**). Expression of IL-6 and IL-8 may further amplify the inflammatory response (**6**) and also in return benefit SARS-CoV-2 infection.

## Material and Methods

### Cell culture

HEK293T cells (Cat. # CRL-3216, ATCC) were cultured in Dulbecco’s modified Eagle’s medium (DMEM, Cat # D5796, Sigma). A549 cells (Cat. # CCL-185, ATCC) were cultured in F12K medium (Cat. # 21127030, Gibco™). Vero E6 cells (Cat. # CRL-1586, ATCC) were cultured in DMEM. HEK293T cells stably expressing ACE2-GFP were previously described *(28)*. A549-ACE2 cells were obtained through BEI Resources, NIH, NIAID (Cat # NR53821). Cell culture medium contained 10% fetal bovine serum (FBS, Cat. # 10437028, Thermo Fisher), penicillin (100 U/ml) /streptomycin (100 μg/ml) (Cat. # MT30002CI, Corning).

### Compounds and antibodies

Recombinant human TNF-α (Cat. # 554618) was purchased from BD. Biosciences. Ribavirin (RIB, Cat. # R0077) was purchased from Tokyo Chemical Industry (TCI). Mycophenolic acid (MPA, Cat. # M3546) was purchased from Sigma-Aldrich. Anti-V5 (Cat. # R960-25), HRP-conjugated anti-V5, and goat HRP-conjugated anti-mouse IgG (H+L) secondary antibody (Cat. # 31430) were purchased from Thermo Fisher Scientific. Anti-GAPDH antibody (Cat. # sc-32233) was purchased from Santa Cruz Biotechnology. Anti-FLAG (Cat. # 2368) antibody, anti-H3 antibody (Cat. # 9715S), and goat HRP-conjugated anti-rabbit IgG antibody (Cat. # 7074) were purchased from Cell Signaling Technology. Anti-IL8 antibody (Cat. # 554717) was purchased from BD. Biosciences.

### Plasmids

pLEX-IMPDH2-V5 vector was picked from the MISSION TRC3 human LentiORF library from Sigma-Aldrich. The pcDNA-FLAG-V5-Nsp10/14/16 vectors were constructed from pDONR223 SARS-CoV-2 Nsp10 (Cat. # 141264, Addgene), Nsp14 (Cat. # 141267, Addgene), and Nsp16 (Cat. # 141269, Addgene) vectors to the pcDNA3.1-3xFLAG-V5-ccdB (Cat. # 87064, Addgene) destination vector using Gateway™ LR Clonase™ II Enzyme Mix (Cat. # 11791020, Invitrogen). pEZY-FLAG-Nsp14 vector was constructed from pDONR223 SARS-CoV-2 Nsp14 vector to the pEZY-FLAG (Cat # 18700, Addgene) destined vector. The pLEX-FLAG-V5 vector was constructed by cloning the FLAG sequence to the pLEX-307 (Cat # 41392, Addgene) vector. The pNF-κB-luciferase vector (PRDII4–luc in the pGL3 vector) was the gift from Dr. Jacob Yount’s lab *(61)*. The pIRES-luciferase vector (Cat. # 219092) was acquired from Agilent Technologies. The pRL-TK Renilla Luciferase vector (Cat. # AF025846) was purchased from Promega.

### Transient transfection

For Nsp14 overexpression, we performed the transient transfection in HEK293T or A549 cells using TurboFect transfection reagents (Cat. # R0531, Thermo Scientific). Briefly, cells were seeded and incubated with the mixture of plasmids with Turbofect (2 µg plasmid DNA / ~3 ×10^5^ cells) for 24 h. The medium was changed, followed by treatment of TNF-α or compounds. For IMPDH2 knockdown, 50 nM siRNA (IMPDH2 assay ID: s7417, sense: 5’-CCAAGAAAAUCACUCUUtt-3’; anti-sense: 5’-UUAAGAGUGAUUUUCUUGGtc-3’, Ambion by Life technologies; non-targeting control: Silencer™ Negative Control No. 4 siRNA, si NT, Cat. # AM4641, Invitrogen) was reversely transfected in HEK293T cells using Lipofectamine™ RNAiMAX Transfection Reagent (Cat. # 13778030, Invitrogen). Cells were kept in culture for 48h and subjected to qPCR analysis for measurement of gene expression.

### Protein immunoblotting

Protein immunoblotting was performed following our previously published protocols *(62, 63)*. Briefly, cells were harvested, washed by PBS, and pelleted. Cell pellets were lysed in RIPA buffer (Cat. #20-188, Millipore) containing protease inhibitor cocktail (Cat. # A32965, Thermo Scientific) on ice, followed by brief sonication to prepare cell lysate. The BCA assay kit (Cat. #23225, Thermo Scientific) was used to quantify the total protein amount in cell lysate, which was boiled in the SDS loading buffer with 5% β-mercaptoethanol (Cat. #60-24-2, Acros Organics). The denatured protein samples were separated by Novex™ WedgeWell™ 4-20% SDS-PAGE Tris-Glycine gel and transferred to PVDF membrane (iBlot™ 2 Transfer Stacks, Invitrogen) using iBlot 2 Dry Blotting System (Cat. # IB21001, Thermo Scientific). The membranes were blocked by 5% milk in PBST and probed by the specific primary antibodies at 4°C overnight, followed by the HRP-conjugated secondary antibodies. The membranes were developed using the Clarity Max ECL substrate (Cat. # 1705062, Bio-Rad).

### Luciferase reporter assays

HEK293T cells were transfected with ISRE or NF-κB luciferase vector along with pRL-TK renilla luciferase vector with or without the indicated vector expressing Nsp14. At 24 h post of transfection, the medium was changed, and cells were treated with 10 ng/ml TNF-α or un-treated for 24h. Cells were lysed using the Dual-Glo® Luciferase Assay System (Cat. #E2920, Promega). Luciferase/renilla signal intensity was detected using Biotek Cytation5 and analyzed by GEN5 software (Biotek).

### Nuclear and cytoplasmic extraction

HEK293T cells were transfected by pcDNA-FLAG-V5-Nsp14 or control vector pLex307-FLAG-V5 for 24h and changed fresh completed DMEM medium for further 24 h culture. Cells (~5 × 10^6^ cells) were collected, washed twice with 1× PBS, and subjected to the nucleus and cytoplasm extraction using NE-PER Nuclear and Cytoplasmic Extraction Reagents (Cat. #78833, Thermo Scientific) following the manufacturer’s instructions and our previous studies *(62)*. Total proteins in the whole-cell lysates from the same number of cells were extracted using 1× RIPA buffer. Extractions from nuclear, cytoplasmic proteins and the total cell lysate proteins were denatured and boiled with 4× LDS sample buffer (Cat. #NP0007, Invitrogen) and subjected to immunoblotting analysis with equal protein loading of extracts (~20 µg/lane). Anti-GAPDH and anti-histone H3 immunoblotting were used as internal controls to determine the cytoplasmic and nuclear fractions.

### Protein co-immunoprecipitation (co-IP)

Protein co-IP assays were performed following the previously published protocol *(62)*. Briefly, protein A/G magnetic beads (Cat. # 88802, Thermo Scientific) and anti-FLAG M2 magnetic beads (Cat. # M8823, Sigma-Aldrich) were washed with 1× RIPA buffer containing protease inhibitor cocktail. Cellular lysates were precleared with the empty magnetic beads for 1 h at 4°C on a 360° tube rocker. The cell lysate was incubated with anti-FLAG M2 magnetic beads for pull-down of FLAG-Nsp14 protein at 4°C overnight with constant rotation. Protein immunocomplexes were washed by RIPA buffer and boiled in SDS loading buffer containing 5% 2-mercaptoethanol, followed by protein immunoblotting. A normal mouse IgG antibody (Cat. # sc-2025, Santa Cruz) was used as the control in parallel.

### Quantitative reverse transcription PCR (RT-qPCR)

RT-qPCR assays were performed following the previously published protocol *(64)*. Total RNAs from harvested cells were extracted using the NucleoSpin RNA extraction kit (Cat. # 740955.250, MACHEREY-NAGEL), and 0.4-1 μg RNA was reversely transcribed using the iScript™ cDNA Synthesis Kit (Cat. # 1708890, Bio-Rad). Real-time qPCR was conducted using the iTaq™ Universal SYBR® GreenSupermix (Cat. # 1727125, Bio-Rad). The PCR reaction was performed on a Bio-Rad CFX connect qPCR machine under the following conditions: 95 °C for 10 m, 50 cycles of 95 °C for 15 s, and 60 °C for 1 m. Relative gene expression was normalized to GAPDH internal control as the 2^-ΔΔCt^ method: 2 ^(ΔCT of targeted gene - ΔCT of GAPDH)^. The following primers were used. IL-4 forward: 5’-GTTCTACAGCCACCATGAGAA-3’, reverse: 5’-CCGTTTCAGGAATCAGATCA-3’; IL-6 forward: 5’-ACTCACCTCTTCAGAACGAATTG-3’, reverse: 5’-CCATCTTTGGAAGGTTCAGGTTG-3’*(30)*; IL-8 forward: 5’-CTTGGCAGCCTTCCTGATTT-3’; reverse: 5’-GGGTGGAAAGGTTTGGAGTATG-3’; Nsp14 forward: 5’-CGGAAACCCAAAGGCTATCA-3’, reverse: 5’-TGTGGGTAGCGTAAGAGTAGAA-3’; IMPDH2 forward: 5′-CTCCCTGGGTACATCGACTT-3′, reverse: 5′-GCCTCTGTGACTGTGTCCAT-3′*(64)*; GAPDH forward: 5′-GCCTCTTGTCTCTTAGATTTGGTC-3′, reverse: 5′-TAGCACTCACCATGTAGTTGAGGT-3′. SARS-CoV-2-TRS-L (N sgRNA forward): CTCTTGTAGATCTGTTCTCTAAACGAAC, SARS-CoV-2-TRS-N (N sgRNA reverse):GGTCCACCAAACGTAATGCG*(65)*

### Viral infection

SARS-CoV-2 strain USA-WA1/2020 was obtained from BEI Resources, NIH, NIAHD (Cat # NR52281) and was plaque purified in Vero E6 cells to identify plaques lacking furin cleavage site mutations. A WT virus plaque was then propagated on Vero E6 cells stably expressing TMPRSS2 (kindly provided by Dr. Shan-Lu Liu, Ohio State University) for 72 h. The virus was aliquoted, flash-frozen in liquid nitrogen, and stored at -80C. The virus stock was titered on Vero E6 cells by TCID50 assay. For infection experiments, the virus was added to cells for 24 h. Cells were then collected by trypsinization and were either lysed with Trizol reagent for RNA extraction or were fixed with 4% paraformaldehyde in PBS for 1 h prior to staining for flow cytometry. Staining was performed with anti-SARS-CoV-2 N (Cat # 40143-MM08, Sino Biological) as described previously *(28, 66)*. Flow cytometry was performed on a FACSCanto II machine (BD Biosciences). Data were analyzed using FlowJo software.

### Human subjects

The lung specimens from deceased COVID-19 patients were obtained from Biobank at Columbia University Irving Medical Center. The control normal lung specimens were the gifts from Jahar Bhattacharya (Columbia University, NY, USA). For paraffin sections, The lungs were fixed with 4% paraformaldehyde (PFA) at 4°C overnight, dehydrated through a series of grade ethanol, and incubated with Histo-Clear (Cat.5989-27-5, National Diagnostics, USA) at room temperature for 2 hours prior to paraffin embedding. 7 μm thick sections were then prepared from the paraffin blocks and mounted on the slides for staining.

### Protein immunofluorescence

Paraffin-embedded lung tissue blocks were baked on the hotplate at 75 °C for 20 min and deparaffinized in xylene. The slides were rehydrated from 100%, 90%, to 70% alcohol, and then to PBS. We performed the antigen unmasking using the retriever (Cat. # 62700-10, Electron Microscopy Sciences) with R-Buffer A pH 6.0 (Cat. # 62706-10, Electron Microscopy Sciences) for 2 h to complete the cycle and cool down. Slides were blocked with 20% normal goat serum (NGS) in PBST for 2 h at room temperature. Slides were incubated with an anti-IL-8 antibody (Cat. # 550419, BD Pharmingen™) in 5% NGS with PBS at 4°C for overnight. Slides were washed with PBST and incubated with Alexa 488 coated goat anti-mouse antibody in 5% NGS/PBS for 2 h at room temperature. Slides were washed with PBST and stained with Hoechst (1:5000 in PBS, Invitrogen). Coverslips were mounted on slides using ProLong Glass Antifade Mountant (Cat. # P36982, Invitrogen) and dried out in the dark overnight. Confocal images were acquired using the ZEISS LSM 700 Upright laser scanning confocal microscope and ZEN imaging software (ZEISS).

### Statistics

Statistical analysis was performed using the GraphPad PRISM. Data are presented as mean ± SD of biological repeats from at least 2 independent experiments. * p<0.05, ** p<0.01, *** p<0.001, or **** p<0.001 indicated the significant difference analyzed by ANOVA or Student’s t-test.

## Acknowledgments

The authors thank Dr. Mark Peeples (Nationwide Children’s Hospital) and Dr. Jianrong Li (The Ohio State University) for kindly providing plaque purified SARS-CoV-2 for viral propagation. We thank Dr. Sheng-Ce Tao (Shanghai Jiao Tong University) for providing Nsp14 cloning plasmid. We also thank Dr. Karin Musier-Forsyth and Dr. Shan-Lu Liu at The Ohio State University for their advice on our studies. This study was funded by NIH research grants R01AI150448, R01DE025447, and R33AI116180 to J.Z., and R03DE029716, R01CA260690 to N.S.

---

**Fig S1.**
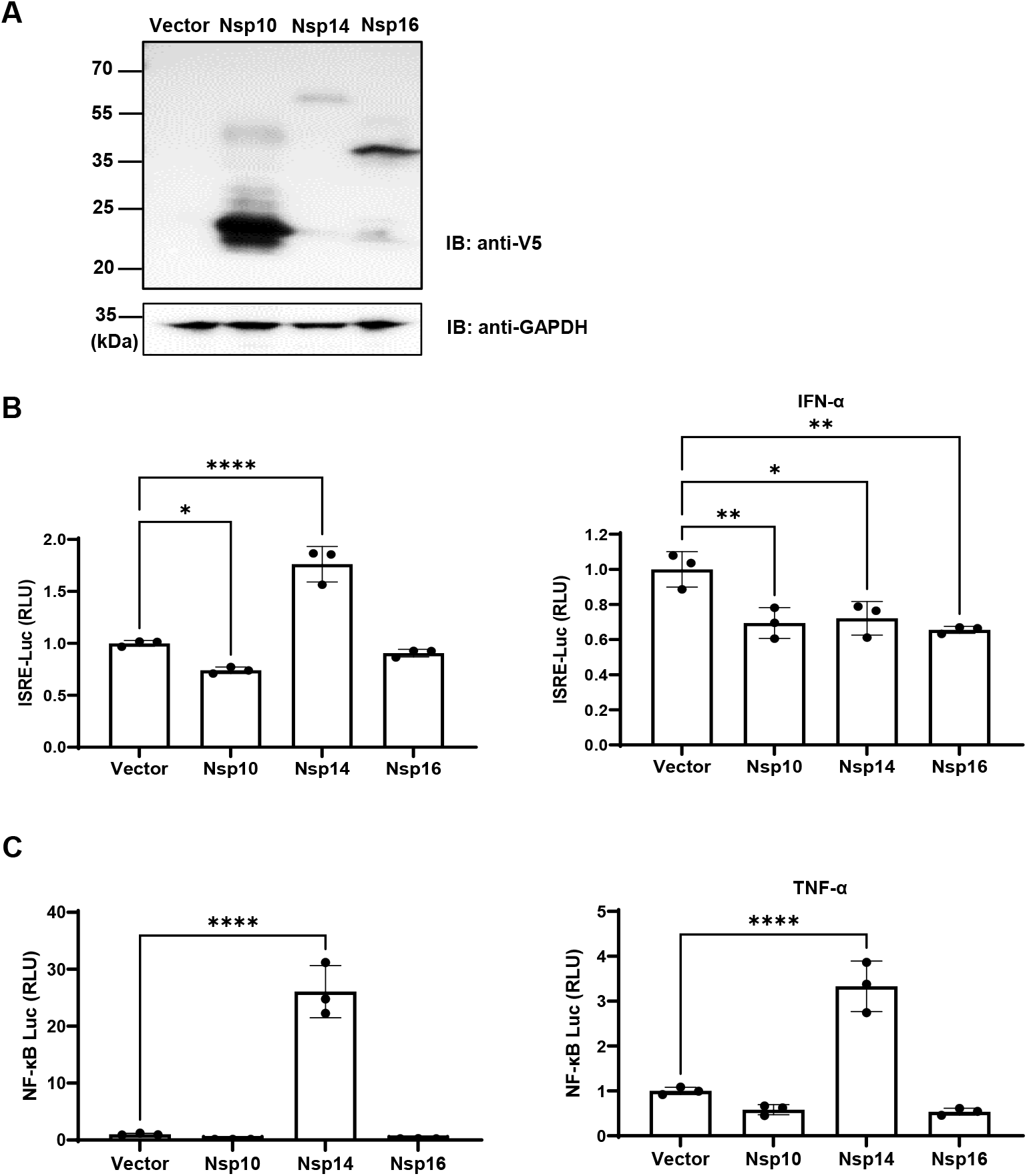
**(A)** HEK293T cells were transiently transfected with the vector expressing V5-FLAG-Nsp10, 14, 16, or empty vector. Protein expression was analyzed by protein immunoblotting. **(B**) HEK293T cells transfected with the vector expressing V5-FLAG-Nsp10, 14, 16, or empty vector along with ISRE-driven firefly luciferase and TK-driven renilla luciferase reporter vectors were un-treated or treated with IFN-α. **(C)** HEK293T cells transfected with the vector expressing V5-FLAG-Nsp10, 14, 16, or empty vector along with NF-κB-driven firefly luciferase and TK-driven renilla luciferase reporter vectors were un-treated or treated with TNF-α. Luciferase activity (firefly/renilla) in these cells was measured and normalized to the empty vector. Results were calculated from 3 technical repeats and presented as mean +/- standard deviation (SD). (* p <0.05; ** p <0.01; *** p <0.001; **** p <0.0001 by unpaired Student’s t-test).

**Fig S2.**
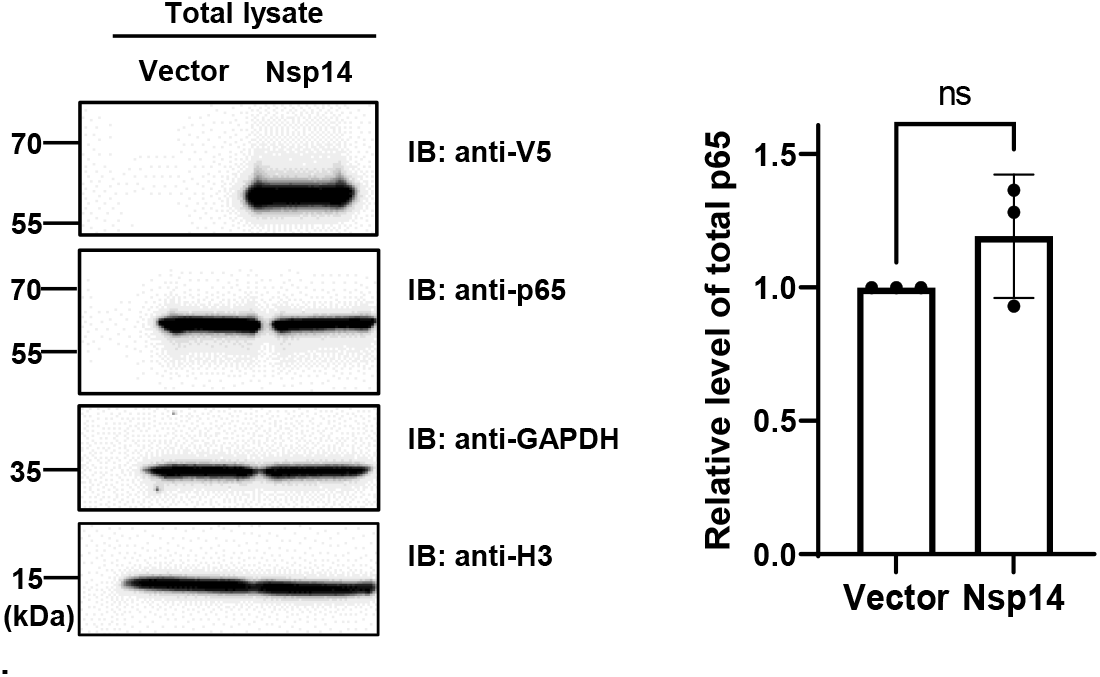
NF-κB p65 in the total lysate of HEK293T cells transfected with the vector expressing V5-FLAG-Nsp14 or empty vector was analyzed by protein immunoblotting. Histone H3 was used as the nuclear marker. The intensity of the p65 protein band was quantified and normalized to the empty vector.

**Fig S3.**
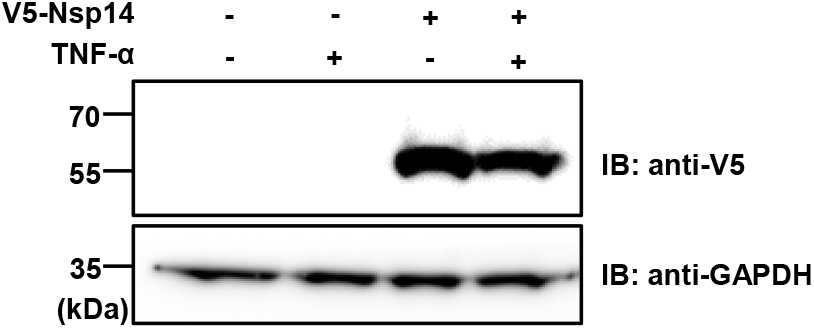
A549 cells were transiently transfected with the vector expressing V5-Nsp14 or empty vector, and treated with or without TNFα. V5-Nsp14 was analyzed by protein immunoblotting.

**Fig S4.**
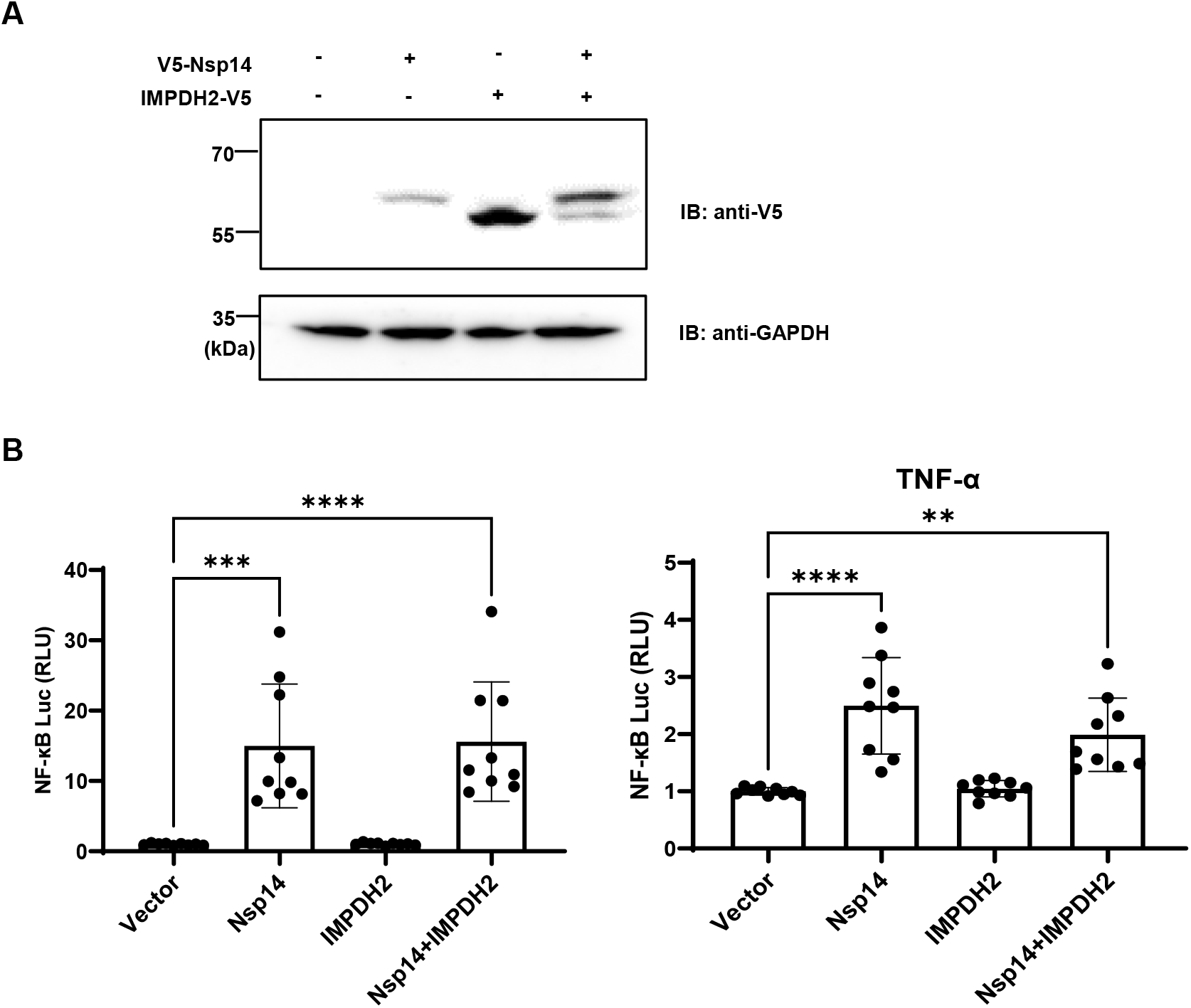
(**A**) HEK293T cells were transiently transfected with the vector expressing V5-Nsp14 or V5-IMPDH2, alone or together. Protein expression was analyzed by protein immunoblotting. (**B**) Cells in (A) further transfected with NF-κB-driven firefly luciferase and TK-driven renilla luciferase reporter vectors were un-treated or treated with TNF-α. Luciferase activity (firefly/renilla) was measured and normalized to the empty vector. Results were calculated from 3 independent experiments and presented as mean +/- standard deviation (SD). (** p < 0.01; *** p <0.001; **** p < 0.0001 by one-way ANOVA and Tukey’s multiple comparison test)

**Fig S5.**
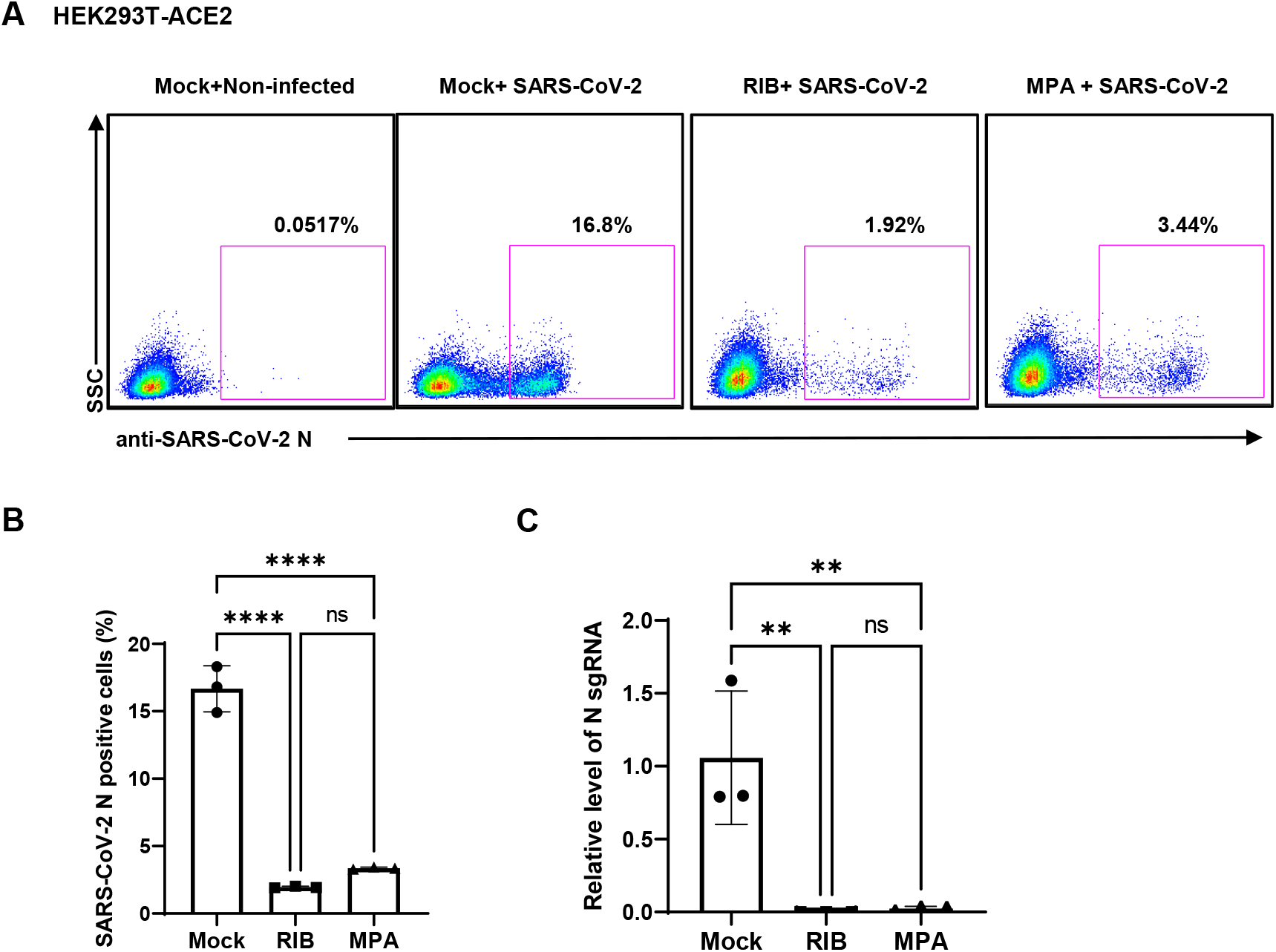
HEK293T-ACE2 cells were treated with RIB (500 µM), or MPA (100 µM), or mock, and infected with SARS-Cov-2 viruses for 24 h. The SARS-CoV-2 infection was detected by intracellular staining of SARS-CoV-2 N protein (**A**). Percentage of SARS-CoV-2 N protein positive cells was calculated (**B**). Cells were harvested for RNA extraction, and N protein sgRNA was analyzed and normalized to the mock treatment (**C**). The results were calculated from 3 technical repeats and presented as mean +/- standard deviation (SD). (** p <0.01; **** p <0.001 by one-way ANOVA and Tukey’s multiple comparison test)

## References

1. F. Robson et al., Coronavirus RNA Proofreading: Molecular Basis and Therapeutic Targeting. Mol Cell 80, 1136–1138 (2020).

2. A. A. Naqvi et al., Insights into SARS-CoV-2 genome, structure, evolution, pathogenesis and therapies: Structural genomics approach. Bba-Mol Basis Dis 1866, (2020).

3. X. Lei et al., Activation and evasion of type I interferon responses by SARS-CoV-2. Nat Commun 11, 3810 (2020).

4. C. K. Yuen et al., SARS-CoV-2 nsp13, nsp14, nsp15 and orf6 function as potent interferon antagonists. Emerg Microbes Infect 9, 1418–1428 (2020).

5. D. Blanco-Melo et al., Imbalanced Host Response to SARS-CoV-2 Drives Development of COVID-19. Cell 181, 1036–1045 e1039 (2020).

6. M. Sa Ribero, N. Jouvenet, M. Dreux, S. Nisole, Interplay between SARS-CoV-2 and the type I interferon response. PLoS Pathog 16, e1008737 (2020).

7. M. Aid et al., Vascular Disease and Thrombosis in SARS-CoV-2-Infected Rhesus Macaques. Cell 183, 1354–1366 e1313 (2020).

8. J. S. Y. Ho et al., TOP1 inhibition therapy protects against SARS-CoV-2-induced lethal inflammation. Cell, (2021).

9. Y. Ma et al., Structural basis and functional analysis of the SARS coronavirus nsp14-nsp10 complex. Proc Natl Acad Sci U S A 112, 9436–9441 (2015).

10. F. Ferron et al., Structural and molecular basis of mismatch correction and ribavirin excision from coronavirus RNA. Proc Natl Acad Sci U S A 115, E162–E171 (2018).

11. E. Minskaia et al., Discovery of an RNA virus 3’->5’ exoribonuclease that is critically involved in coronavirus RNA synthesis. Proc Natl Acad Sci U S A 103, 5108–5113 (2006).

12. M. Bouvet et al., RNA 3’-end mismatch excision by the severe acute respiratory syndrome coronavirus nonstructural protein nsp10/nsp14 exoribonuclease complex. Proc Natl Acad Sci U S A 109, 9372–9377 (2012).

13. N. S. Ogando et al., The Enzymatic Activity of the nsp14 Exoribonuclease Is Critical for Replication of MERS-CoV and SARS-CoV-2. J Virol 94, (2020).

14. N. H. Moeller et al., Structure and dynamics of SARS-CoV-2 proofreading exoribonuclease ExoN. bioRxiv, (2021).

15. Y. Chen et al., Functional screen reveals SARS coronavirus nonstructural protein nsp14 as a novel cap N7 methyltransferase. Proc Natl Acad Sci U S A 106, 3484–3489 (2009).

16. M. Bouvet et al., In vitro reconstitution of SARS-coronavirus mRNA cap methylation. PLoS Pathog 6, e1000863 (2010).

17. Y. Chen et al., Biochemical and structural insights into the mechanisms of SARS coronavirus RNA ribose 2’-O-methylation by nsp16/nsp10 protein complex. PLoS Pathog 7, e1002294 (2011).

18. E. Decroly, F. Ferron, J. Lescar, B. Canard, Conventional and unconventional mechanisms for capping viral mRNA. Nat Rev Microbiol 10, 51–65 (2011).

19. Z. A. Jaafar, J. S. Kieft, Viral RNA structure-based strategies to manipulate translation. Nat Rev Microbiol 17, 110–123 (2019).

20. E. Jan, I. Mohr, D. Walsh, A Cap-to-Tail Guide to mRNA Translation Strategies in Virus-Infected Cells. Annu Rev Virol 3, 283–307 (2016).

21. M. Becares et al., Mutagenesis of Coronavirus nsp14 Reveals Its Potential Role in Modulation of the Innate Immune Response. J Virol 90, 5399–5414 (2016).

22. J. Gribble et al., The coronavirus proofreading exoribonuclease mediates extensive viral recombination. PLoS Pathog 17, e1009226 (2021).

23. D. E. Gordon et al., A SARS-CoV-2 protein interaction map reveals targets for drug repurposing. Nature 583, 459–468 (2020).

24. L. X. Liao et al., Highly selective inhibition of IMPDH2 provides the basis of antineuroinflammation therapy. Proc Natl Acad Sci U S A 114, E5986–E5994 (2017).

25. T. Liu, L. Zhang, D. Joo, S. C. Sun, NF-kappaB signaling in inflammation. Signal Transduct Target Ther 2, (2017).

26. C. Grassl, B. Luckow, D. Schlondorff, U. Dendorfer, Transcriptional regulation of the interleukin-6 gene in mesangial cells. J Am Soc Nephrol 10, 1466–1477 (1999).

27. V. Bezzerri et al., Mapping the transcriptional machinery of the IL-8 gene in human bronchial epithelial cells. J Immunol 187, 6069–6081 (2011).

28. G. Shi et al., Opposing activities of IFITM proteins in SARS-CoV-2 infection. Embo J 40, e106501 (2021).

29. L. Leng et al., Pathological features of COVID-19-associated lung injury: a preliminary proteomics report based on clinical samples. Signal Transduct Target Ther 5, 240 (2020).

30. J. C. Melms et al., A molecular single-cell lung atlas of lethal COVID-19. Nature, (2021).

31. L. Hedstrom, IMP dehydrogenase: structure, mechanism, and inhibition. Chem Rev 109, 2903–2928 (2009).

32. S. Zhou, R. Liu, B. M. Baroudy, B. A. Malcolm, G. R. Reyes, The effect of ribavirin and IMPDH inhibitors on hepatitis C virus subgenomic replicon RNA. Virology 310, 333–342 (2003).

33. P. Leyssen, J. Balzarini, E. De Clercq, J. Neyts, The predominant mechanism by which ribavirin exerts its antiviral activity in vitro against flaviviruses and paramyxoviruses is mediated by inhibition of IMP dehydrogenase. J Virol 79, 1943–1947 (2005).

34. M. D. Sintchak et al., Structure and mechanism of inosine monophosphate dehydrogenase in complex with the immunosuppressant mycophenolic acid. Cell 85, 921–930 (1996).

35. C. W. Yang et al., Targeting Coronaviral Replication and Cellular JAK2 Mediated Dominant NF-kappaB Activation for Comprehensive and Ultimate Inhibition of Coronaviral Activity. Sci Rep 7, 4105 (2017).

36. N. Hemmat et al., The roles of signaling pathways in SARS-CoV-2 infection; lessons learned from SARS-CoV and MERS-CoV. Arch Virol 166, 675–696 (2021).

37. X. Yin et al., MDA5 Governs the Innate Immune Response to SARS-CoV-2 in Lung Epithelial Cells. Cell Rep 34, 108628 (2021).

38. M. M. Rahman, G. McFadden, Modulation of NF-kappaB signalling by microbial pathogens. Nat Rev Microbiol 9, 291–306 (2011).

39. J. L. Forbester, I. R. Humphreys, Genetic influences on viral-induced cytokine responses in the lung. Mucosal Immunol 14, 14–25 (2021).

40. R. Alon et al., Leukocyte trafficking to the lungs and beyond: lessons from influenza for COVID-19. Nat Rev Immunol 21, 49–64 (2021).

41. C. M. Ziegler et al., A Proteomics Survey of Junin Virus Interactions with Human Proteins Reveals Host Factors Required for Arenavirus Replication. J Virol 92, (2018).

42. W. Dang et al., Inhibition of Calcineurin or IMP Dehydrogenase Exerts Moderate to Potent Antiviral Activity against Norovirus Replication. Antimicrob Agents Chemother 61, (2017).

43. W. P. Hofmann, E. Herrmann, C. Sarrazin, S. Zeuzem, Ribavirin mode of action in chronic hepatitis C: from clinical use back to molecular mechanisms. Liver Int 28, 1332–1343 (2008).

44. L. Quemeneur et al., Differential control of cell cycle, proliferation, and survival of primary T lymphocytes by purine and pyrimidine nucleotides. J Immunol 170, 4986–4995 (2003).

45. Q. N. Shu, V. Nair, Inosine monophosphate dehydrogenase (IMPDH) as a target in drug discovery. Med Res Rev 28, 219–232 (2008).

46. J. Toubiana et al., IMPDHII protein inhibits Toll-like receptor 2-mediated activation of NF-kappaB. J Biol Chem 286, 23319–23333 (2011).

47. S. Kofuji et al., IMP dehydrogenase-2 drives aberrant nucleolar activity and promotes tumorigenesis in glioblastoma. Nat Cell Biol 21, 1003–1014 (2019).

48. S. Kofuji, A. T. Sasaki, GTP metabolic reprogramming by IMPDH2: unlocking cancer cells’ fuelling mechanism. J Biochem 168, 319–328 (2020).

49. Q. Zhang et al., The role of IMP dehydrogenase 2 in Inauhzin-induced ribosomal stress. Elife 3, (2014).

50. S. Mannava et al., Direct role of nucleotide metabolism in C-MYC-dependent proliferation of melanoma cells. Cell Cycle 7, 2392–2400 (2008).

51. J. Zhang et al., A systemic and molecular study of subcellular localization of SARS-CoV-2 proteins. Signal Transduct Target Ther 5, 269 (2020).

52. J. M. Meyers et al., The proximal proteome of 17 SARS-CoV-2 proteins links to disrupted antiviral signaling and host translation. bioRxiv, (2021).

53. B. R. Lane et al., Interleukin-8 stimulates human immunodeficiency virus type 1 replication and is a potential new target for antiretroviral therapy. J Virol 75, 8195–8202 (2001).

54. W. C. Chen et al., HCV NS5A Up-Regulates COX-2 Expression via IL-8-Mediated Activation of the ERK/JNK MAPK Pathway. Plos One 10, e0133264 (2015).

55. N. Mukaida, Pathophysiological roles of interleukin-8/CXCL8 in pulmonary diseases. Am J Physiol Lung Cell Mol Physiol 284, L566–577 (2003).

56. T. Murayama et al., Enhancement human cytomegalovirus replication in a human lung fibroblast cell line by interleukin-8. J Virol 68, 7582–7585 (1994).

57. J. Hiscott, H. Kwon, P. Genin, Hostile takeovers: viral appropriation of the NF-kappaB pathway. J Clin Invest 107, 143–151 (2001).

58. I. F. Hung et al., Triple combination of interferon beta-1b, lopinavir-ritonavir, and ribavirin in the treatment of patients admitted to hospital with COVID-19: an open-label, randomised, phase 2 trial. Lancet 395, 1695–1704 (2020).

59. W. Wan et al., High-Throughput Screening of an FDA-Approved Drug Library Identifies Inhibitors against Arenaviruses and SARS-CoV-2. ACS Infect Dis, (2020).

60. Y. Han et al., Identification of SARS-CoV-2 inhibitors using lung and colonic organoids. Nature 589, 270–275 (2021).

61. E. Prinarakis, E. Chantzoura, D. Thanos, G. Spyrou, S-glutathionylation of IRF3 regulates IRF3-CBP interaction and activation of the IFN beta pathway. Embo J 27, 865–875 (2008).

62. D. Zhou et al., Inhibition of Polo-like kinase 1 (PLK1) facilitates the elimination of HIV-1 viral reservoirs in CD4(+) T cells ex vivo. Sci Adv 6, eaba1941 (2020).

63. W. Kong et al., Nucleolar protein NOP2/NSUN1 suppresses HIV-1 transcription and promotes viral latency by competing with Tat for TAR binding and methylation. PLoS Pathog 16, e1008430 (2020).

64. F. Huang et al., Inosine Monophosphate Dehydrogenase Dependence in a Subset of Small Cell Lung Cancers. Cell Metab 28, 369–382 e365 (2018).

65. L. Yang et al., A Human Pluripotent Stem Cell-based Platform to Study SARS-CoV-2 Tropism and Model Virus Infection in Human Cells and Organoids. Cell Stem Cell 27, 125–136 e127 (2020).

66. R. C. Larue et al., Rationally Designed ACE2-Derived Peptides Inhibit SARS-CoV-2. Bioconjug Chem 32, 215–223 (2021).

